# Productivity and resilience of Chinook salmon compromised by ‘Mixed-Maturation’ fisheries in marine waters

**DOI:** 10.1101/2024.04.25.591098

**Authors:** Nick Gayeski, Devin Swanson, Misty MacDuffee, Andrew Rosenberger

## Abstract

Most Chinook salmon (*Oncorhynchus tshawytscha*) in the northeast Pacific Ocean are harvested in mixed-stock marine fisheries. Here, multiple populations with varying abundance and productivities are encountered. In addition, many of these fisheries generally encounter both mature and immature Chinook. Hence, these fisheries are better described as mixed-stock and “mixed-maturation” (MM) fisheries. Harvest of immature fish can skew the age composition of Chinook populations towards younger, and hence smaller, individuals. Older Chinook are generally larger and contribute disproportionately to the productivity of their populations. We developed an individual-based demographic-genetic model of ocean-type Chinook to evaluate the effects of fisheries that harvest immature Chinook. We then compared those effects to terminal fisheries that harvest only mature fish. Our model provides the ability to assess the benefits of terminal Chinook fisheries to both landed catch and Chinook rebuilding. Recovered populations show a more archetypal age– and sex-structure than contemporary ocean-type Chinook subject to marine mixed-maturation fisheries. In our modeled scenarios of mixed-maturation fisheries, we found that immature Chinook can comprise up to 59% of the total numbers of fish caught, and 47% of the total weight of the catch. If instead, these Chinook were not harvested until they mature and reach terminal fisheries, they would contribute greater biomass to landed catches. These terminal fisheries allow a higher percentage of larger, older Chinook to escape, and would increase the fecundity and productivities of their populations. The benefits of terminal fisheries would accrue to fishers, sustainable wild harvesting, wildlife, and the rebuilding of depleted Chinook runs.

## Introduction

A number of wild Pacific salmon species and populations have declined in adult body size over recent decades. Such observations in Chinook salmon (*O. tshawytscha*) have been documented since the 1920s. Chinook salmon have displayed marked declines in size-at-age and declines in the proportion of older, larger individuals, particularly females, in many populations across their range in the eastern Pacific and watersheds from the Pacific Northwest to Alaska [1–12].

A number of likely factors affecting salmon body size over this time have been identified, including marine environmental conditions, interactions with large numbers of hatchery salmon in marine waters [13–15], adverse genetic and ecological interactions with hatchery salmon, and the direct and indirect effects of fisheries that impose selection [8, 9,16–20]. While there have undoubtedly been major changes in the marine ecosystems of the northeastern Pacific in which Chinook populations rear for the majority of their lives, and many of these changes are likely to adversely affect the marine growth of Chinook, such processes do not rule out a significant role for fisheries induced demographic change and/or selection (FIS) (e.g., [18, 21, 22]).

Of particular relevance to ocean-type Chinook salmon populations originating in British Columbia (BC) and the southern US (Washington, Oregon and California, including the lower and middle Columbia River) are coastal mixed-stock fisheries that occur in territorial waters (3 nautical miles) of provincial and state coasts and in the Pacific Coast EEZs of Canada and the US. These fisheries are managed pursuant to the Pacific Salmon Treaty (PST), and domestically through the US Pacific Fisheries Management Council (PFMC) Salmon Plan and Canada’s Integrated Fisheries Management Planning (IFMP) process. The majority of ocean-type Chinook (also called Fall Chinook [23, 24]) from these populations migrate north as sub-yearlings to rear inside of the continental shelf in the coastal marine waters of BC and Southeast Alaska (SEAK) before attaining their maturation size and age and migrating back to their natal rivers. These immature Chinook are subject to harvest in marine troll and net fisheries conducted in SEAK and BC that are managed under the Aggregate Abundance-Based Management (AABM) regime of the PST. Hence, these coastal mixed-stock fisheries are also “mixed-maturation (MM) marine fisheries.” Fall Chinook bound for rivers in the southern US and BC that escape the AABM fisheries are then subject to one or more state, tribal or provincial fisheries managed pursuant to the PFMC or the IFMP Salmon Plans. As a result of this life-history and migratory pattern, immature ocean-type Chinook are vulnerable to capture in coastal mixed-stock fisheries at similar rates as mature individuals.

### Direct and indirect age overfishing

Harvest of large numbers of immature salmon, i.e., individuals captured one or more years before they have completed their marine growth and become mature adults, results in indirect age-overfishing and consequent growth-overfishing (lowered productivity and yield per recruit). Indirect age– and/or size-overfishing is a distinct phenomenon from direct age-overfishing.

Direct age-overfishing occurs when older, typically larger individuals are targeted by harvesters through gear selectivity and/or fishing at times or in areas where older, larger fish are known to occur in numbers disproportionate to their abundance in the total harvestable population. This directly reduces the proportion of older, larger fish in the remaining unharvested population, especially the spawning population. This can also select for an earlier age at maturity either by selecting on the growth rate or on the maturation size, or both. Indirect age– (and thus size-) overfishing, by contrast, reduces the proportion and total abundance of older, larger adults in the harvestable and the spawning population by removing individuals one or more years before they are scheduled to mature when they are at smaller sizes.

Independent of ecological conditions in marine waters that may result in adverse growth conditions, age– and growth-overfishing (direct or indirect) is a result of fishing mortality alone. The effect of intensive and age– and size-selective harvesting on downsizing body size is well documented in the fisheries literature [25–32], as life-history theory predicts that increased mortality in older age and size classes generally favors earlier sexual maturation at smaller size [27, 29, 33]. Indirect age over-fishing is less likely to occur in terminal (e.g., river) or near-terminal (e.g., estuary) fisheries as all, or a majority of, Chinook present are returning mature spawners, (although direct size over-fishing may still occur if the gear and/or the fishers differentially select for larger fish [34]). Further, indirect age-overfishing is less likely than direct age-overfishing to select for faster growth rates (and thus younger maturation age) because when immature individuals are at sizes vulnerable to the fishing gear there is generally no benefit to faster growth, as larger fish are likely to be equally or more vulnerable to fishing gear.

To date, the potential selective and demographic effects of age-overfishing in coastal mixed-maturation Chinook fisheries have not been closely analyzed, though the potential for adverse selective and/or demographic consequences has been acknowledged [2, 8, 18, 25, 35]. We hypothesize that (indirect) age-overfishing in mixed maturation (MM) fisheries results in changes in the demographics of affected Chinook populations, selecting for younger age-at-maturity as a result of the reduction in fitness of immature Chinook genetically programmed to mature at older ages. That is, genes that select for older ages at maturity are reduced in frequency in the population as immature individuals are removed from the population. As a result of adverse impacts on the fitness of older, larger Chinook, indirect age overfishing shifts the age structure of the spawning population toward a younger mean age, with resultant reductions in mean body size, thereby reducing catch biomass and the productivity of the spawning population. Reductions in the proportion and size of older females in particular are likely to have adverse consequences for population productivity, as fecundity is related to the length of female Chinook salmon [6, 12, 36], as well as having adverse impacts on the value of the landed catch (e.g. [37]). If correct, MM fisheries in general are likely to have adverse impacts on the productivity and resilience of extant Chinook populations in comparison to terminal/near-terminal fisheries. This affects the potential for recovery of depressed Chinook populations in BC and the Southern US. In addition, such loss of productivity and resilience is likely to inhibit future returns of harvestable Chinook populations to terminal and near-terminal areas that may provide long-term, sustainable fisheries that support small-scale local fishing economies and communities.

To investigate the potential of MM fisheries to effect harvest-induced change, we developed an individual-based demographic-genetic model [21, 38, 39] based on the life-history of ocean-type Chinook. We then applied the model to evaluate MM harvest of immature and maturing individuals relative to a terminal fishery harvesting only mature adults. An individual-based model simulates demographic and selected genetic processes of each individual in the population, and is thus capable of considering individual variation in demographic and relevant genetic characteristics within size– and/or age-classes. This approach is particularly valuable for addressing issues related to fisheries induced demographic change and selection. To facilitate informative modeling of different fishing gears– particularly troll and gill net fisheries – we parameterized our model based on fork length.

Our primary focus is on the demographic effects of the various harvest regimes. In particular, we are interested in changes in the age-and size-structure of the population due to the effects of harvest mortality on the fitness of the phenotypes produced by the suite of available genotypes that control daily growth rates and maturation lengths. To provide a clear focus on the contrasting effects of MM and terminal fisheries, we conduct our harvest simulation on a single population. Hence, we do not address the additional issues that surround the effects of harvest in mixed stock marine fisheries.

We first describe the basic features of the model and then report on the results of six harvest simulation scenarios, ranging from typical coastal MM Chinook fisheries to terminal fisheries. We evaluate harvest impacts on size– and-age-of maturity of males and females, the proportions of sex and age class in the annual adult return, the proportions of sex and age class on the spawning grounds, and how this relates to reproductive potential. We refer to the model as an “individual-based Chinook demographic-genetic model (IBCDM) for length-based ocean-type life-history”.

## Methods

### Development of the Individual-based Chinook Demographic-genetic Model (IBCDM)

#### General model features

Our model was derived in large part from the IBSEM model of [39] and parameterized to reflect the life-history of ocean-type Chinook salmon. Juvenile ocean-type Chinook become smolts as sub-yearlings, migrating seaward within several weeks or months of emerging from the spawning gravel [23, 24, 40]. We also incorporated some of the model approaches developed by [41] to evaluate fisheries selectivity of an in-river gillnet fishery for Yukon River Chinook. The model was written in C++. For brevity, in the rest of this paper we refer to our model either as IBCDM or simply as “the model”.

Our focus is on comparing mixed-maturation (MM) marine and terminal fisheries and not on mixed-stock fisheries, *per se*. We therefore model a single population with an ocean-type life-history and a size– and age-structure likely to be representative of a 19^th^ century population prior to the development of intensive coastal mixed-stock and MM fisheries. Most importantly, the model population contains a higher proportion of older individuals (ages 5 and 6) than the majority of ocean-type Chinook populations in the latter half of the 20^th^ century and the first 2 decades of the 21^st^ century. The model population is based on a combination of data for ocean-type Chinook salmon from the Fraser River (BC) from the mid-1960s [42] and the Hanford Reach Upriver Bright population in the middle Columbia River provided by [43], who analyzed several data sources between 1920 and 1990. This data was used to build a model we believe is generally representative of an ocean-type Chinook population that may have existed prior to modern fisheries occurring in the Columbia River mainstem, Georgia Strait and the lower Fraser River by the 1920s [3, 23]. Consequently, similar to [41], our model population is best viewed as representing a generic historical ocean-type Chinook population. The model has the flexibility to be parameterized with stock-specific data when that is of interest. For the objectives of this paper, the generic model is appropriate.

The model population consists of the following stages: eggs, parr (post-emergent fry), smolts, and post-smolts of ages 1 to 6 (Supplementary File S2, Table A). Males and females are modeled separately. Males mature between the ages of 2 (ocean age 1) and the maximum age of 6 (ocean age 5); females between the ages of 3 (ocean age 2) and 6 (ocean age 5). Spawning is modeled as occurring on November 1 of each model year, and fry (called “parr”) are modeled as emerging from the gravel and completing yolk absorption on May 1 of the following calendar year. All parr are assumed to have a common size (weight and fork length) at the time of emergence.

Genetics control the probability of maturing at an adult size within length intervals reflective of the growth trajectories of the majority of males and females at each possible age of maturity. Size at maturity is greater than 350 millimeters (mm) fork length (FL) for males and greater than 520 mm for females. Genetic control is parameterized following the approach of [39] and [44]. The general features are described in the **Genetics** section below. Additional details are given in Supplementary File S1.

Both growth and survival are modeled from the parr stage forward. Ricker-type density dependence is included and applies to survival of newly emerged parr from May 1 to smolt at July 1 (60 days) at which time smolts migrate to marine waters. Growth and density-independent survival of smolts is calculated to November 1 of the year of migration (123 days) at which time smolts are referred to as “age 1”, the age they will attain if they survive to their next birthday on May 1 of the following calendar year. Growth and survival from the smolt stage forward is modeled using the joint growth-and-mortality model of McGurk [45]. Parr are assumed to grow in accordance with the growth rate component of the McGurk model but survival to the smolt stage is subject to the size-independent Ricker density-dependent survival equation. From age-1 forward growth, survival, potential harvest and maturation occur at annual time steps. Harvest occurs immediately prior to adult spawning. Ricker-type juvenile density dependence, and growth and post-smolt density independent survival are described in Supplement File S1, A4.

We parameterized the model so that it results in an adult population with a Ricker spawner recruit relationship with an alpha value of 5.24, a beta value of 604, and a deterministic equilibrium adult population size of 1000 (Supplementary File S2, Table K). Fig 1 shows the general life cycle of the population. The model life-stages and related parameters and values are shown in Supplementary File S2 (Tables A, B, and C).

**Fig 1.**
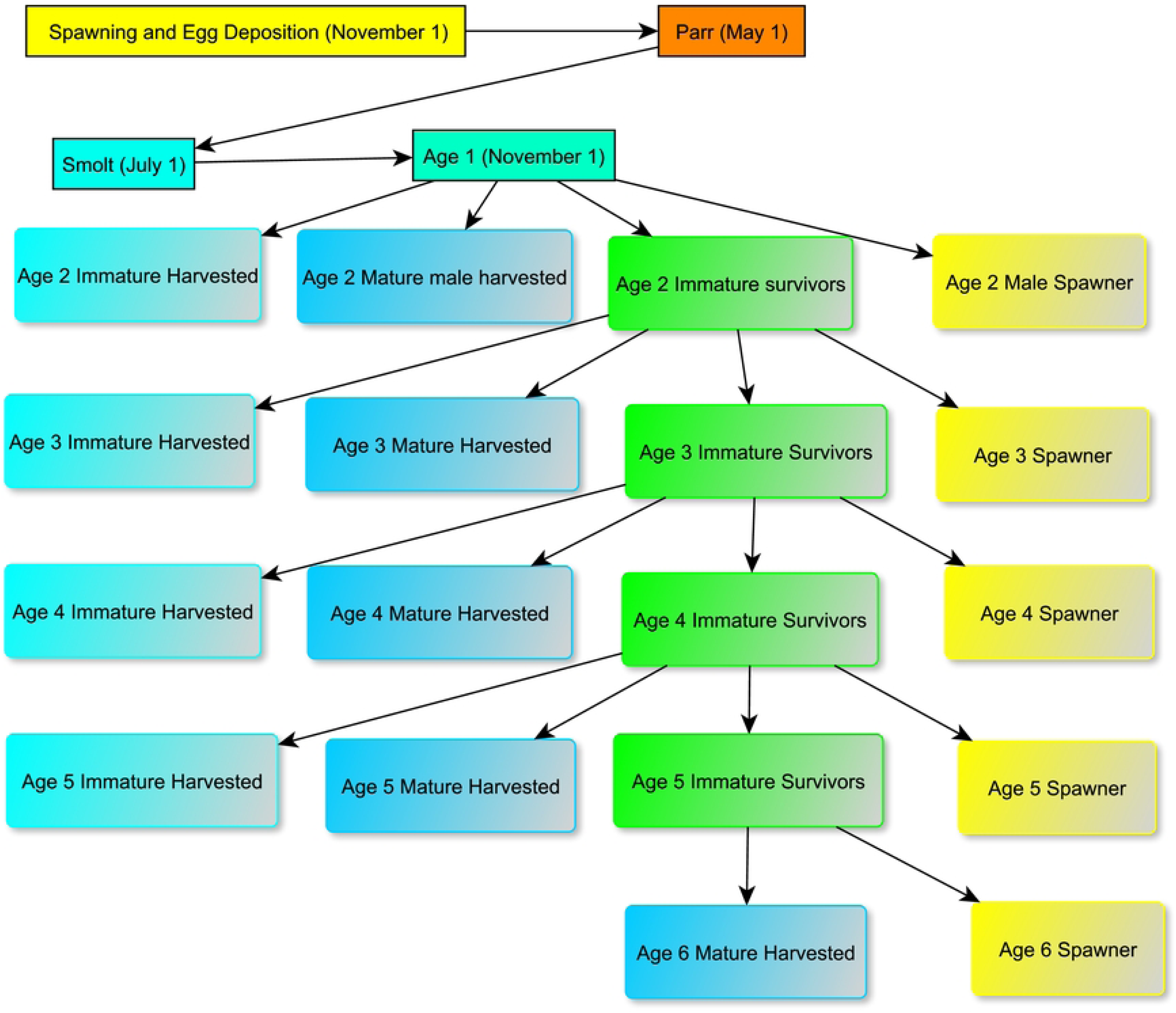
Generalized life history of ocean-type Chinook salmon.

#### Model logic

The model is initialized as a colonization process by a spawning population of 508 males ages 2 to 6, and 492 females ages 3 to 6. The proportions of ages in each sex are chosen to achieve a target adult equilibrium population of approximately 1000 individuals, a sex ratio of approximately 1:1 and target age/sex proportions. Since the model is length– and not age-based, we refer to the initial spawning population by length-intervals associated with the dominant maturation age within each interval (abbreviated L2, L3, etc.), as shown below in Fig 2. At initialization these can be thought of as equal to the mean fork length in millimeters of the maturation age of each sex, listed in Table 1 and Supplementary File S2, Table C.

**Fig 2.**
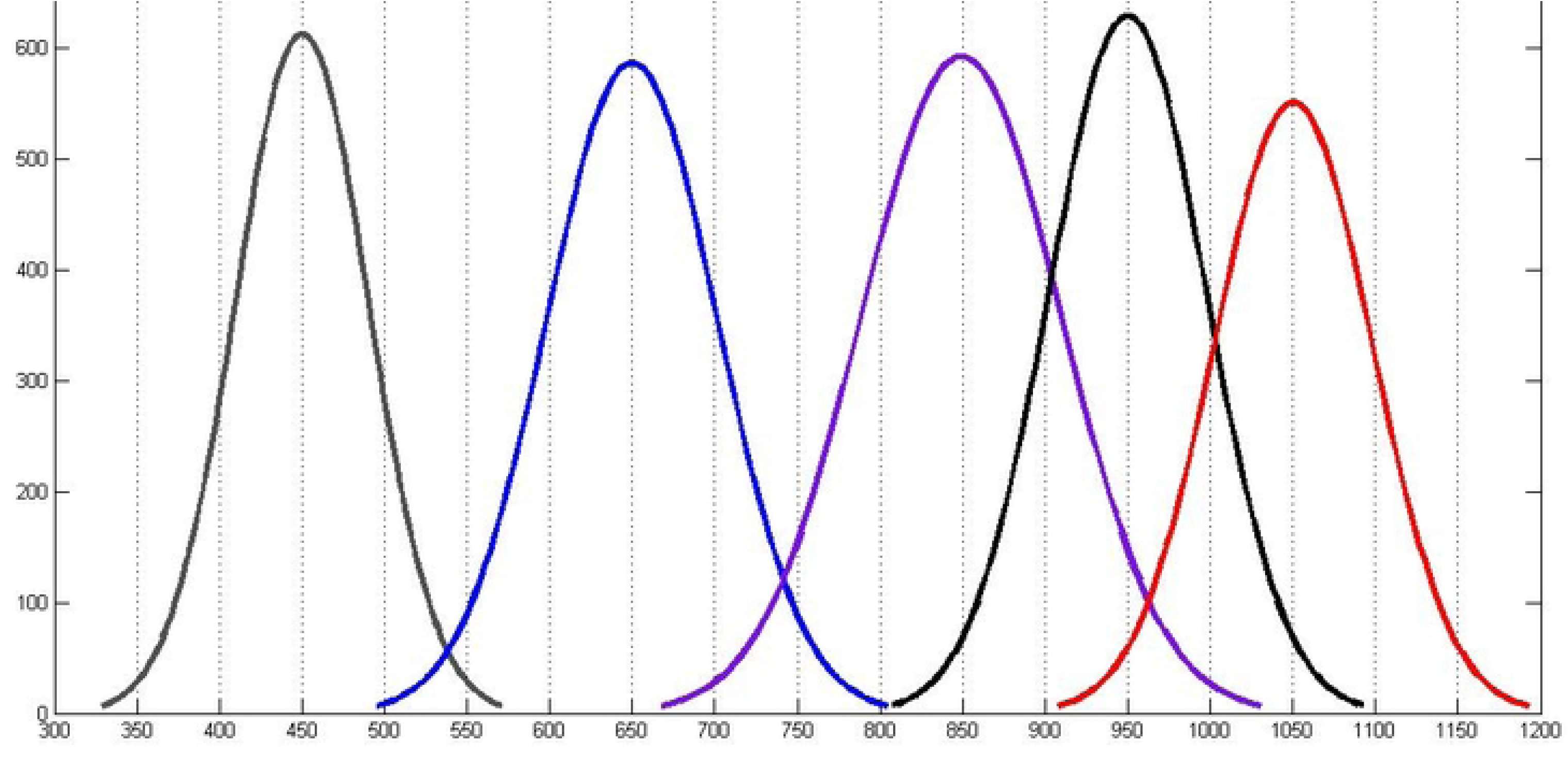
Maturation lengths in mm at ages 2 to 6 from 10000 simulated random values drawn from each of the five possible bivariate-normal distributions of daily growth rates and maturation lengths. age groups L2 to L6 from left to right.

**Table 1.** Mean age-specific lengths and weights (standard deviation) of the ocean-type Chinook model population pre 20^th^ century.

|  | Age 2 | Age 3 | Age 4 | Age 5 | Age 6 |
| --- | --- | --- | --- | --- | --- |
| Mean Fork Length,<br>mm (inches) | 450 (18) | 650 (26) | 850<br>(33.5) | 950 (37.4) | 1050<br>(41.3) |
| Mean Weight, grams<br>(lbs.) | 1020 (2.25) | 3283 (7.24) | 7712<br>(17.0) | 10984<br>(24.2) | 15094<br>(33.3) |

Due to random growth of individuals and size-dependent density independent post-smolt mortality (described in Supplement File S1, and **Post-smolt growth and survival** below), the unfished equilibrium abundance will vary around the target of 500 male and 500 female spawners, but will rarely be exactly these values. We evaluated the time it takes to attain a stochastic equilibrium by running the model without harvest for 1000 time-steps (years) and examining the distributions of spawner sexes and ages at several 25-year (approximately 6 generation) intervals and at the final year of each 25-year period. We determined that the model achieves its stochastic equilibrium by year 100. Data for a 1000-year non-harvest simulation is provided in Supplement File S4.

In the simulations reported below, we run the model for 100 years without harvest, allowing the population to achieve a stochastic equilibrium following the initial colonization. We then run the selected harvest scenarios for 25 years, followed by 25 years of no harvest. We chose 25 years for the harvest period as a reasonable length of time for harvest to impact the demographics of the population in a time frame relevant to short-term management interests, but short enough that fisheries-induced evolution is not likely to occur. Similarly, we chose 25 years for the post-harvest evaluation period as a time frame relevant to a management evaluation of the ability of the harvested population to respond demographically to the cessation of harvest, assuming that no fishery-induced selection occurs during the harvest period. This provides a realistic measure of the demographic impact of the 25-year harvest regime on the population and the population’s ability to recover from those impacts.

#### Genetics

Individuals are diploid. Daily growth rate and maturation size (weight) is defined by a single set of 20 genes. Length is determined from weight following an allometric length-weight equation (L = aW^b^). For computational efficiency sex is randomly assigned at the smolt stage with an equal (mean) sex ratio. Individuals inherit a maternal and a paternal allele at each locus.

We follow [39] in employing an additive quantitative genetics model. The genotypic effect (breeding value) is determined using a simple additive model with exponentially declining weights, such that there are a small number of genes (loci) which have relatively large effect on daily growth rate and size-at-maturity, and the remaining genes having exponentially smaller influence as shown in Supplement File S1, Table A and Fig A.

Maturation age and size (weight, WMat) are determined by quantitative genetic parameters controlling the daily growth rate in weight and the weight– (and hence length-) at-maturity assigned to offspring. Maturation is determined by the “total allelic (genotypic) value” measured across the 20 bi-allelic (0, 1) loci, which is equal to the individual’s breeding value. Each fertilized egg inherits one allele (0 or 1) with equal probability from each parent at each of the 20 loci, by successively choosing at random one parental allele from each locus using a uniform distribution on [0,1]. Following [39] the “allelic value” of each locus is determined by multiplying the genotypic value of each locus (0, 0.5, or 1) by a locus-specific weight. The weights decline exponentially from a maximum value at locus 1 to a minimum at locus 20. (Supplement File S1, Table A and Fig A.) The sum of all weighted allelic values (the sum of “1” alleles weighted by the locus-specific weight of influence, [46]) constitutes the “total allelic value” of an individual’s genotype. Following [39], we name this total genetic value sPhi (S^φ^). The values of S^φ^ range from a minimum of 0 (when all 20 loci are homozygous for the “0” allele) to a maximum of 1 (when all 20 loci are homozygous for the “1” allele).

Each S^φ^ value within a specified range (e.g., >0.2 and <=0.4) then determines the probability of an individual maturing at one of three length intervals by virtue of inheriting a daily growth rate (GR) in weight (g/g body weight) and a target maturation weight (WMat) and length (LMat) from a bivariate normal distribution with mean growth rate (GRx) and mean maturation weight (WxMat). Maturation weight is then converted to maturation length (LxMat) via the deterministic allometric length-weight equation (Supplemental File S2, equation 1.2). LxMat is equal to the maturation length that an individual with mean growth rate GRx would attain at age x, for x = 2 to 6 for males and 3 to 6 for females. Thus, we model the genetic determination of the mean GR and WMat values assigned to offspring as canalized as an evolutionary response to the environmental variability experienced by the founder population during its evolutionary history. This constitutes a form of “diversification bet-hedging” against unpredictable environmental variability that can fundamentally affect the fitness of different maturation ages [47–51].

Additionally, when desired, phenotypic plasticity in growth rate and target maturation weight and length can be allowed to occur based on stochasticity in individual daily growth and annual survival rates that may be experienced in the post-smolt marine environment. This feature is described in the Supplement File S1, **Incorporating environmental stochasticity in maturation weight and length.**

After mating and spawning occur, parr emerge, undergo density-dependent survival to the smolt stage and sex is assigned to each smolt, daily growth rate, maturation weight, and length is determined according to each individual’s total allelic value as described above and in Supplement File S1, Section A2.

#### Spawning and mating

The mean fecundity of a female (mf) is determined by its fork length [36] based on the Pearson equation used by [41]. The actual number of eggs is determined from a normal distribution with mean mf and a standard deviation (mfsd) chosen to achieve an appropriate random variation in fecundity at each female age.

Mating is modeled as size-selective following the approach of [41]. Each female spawner is chosen at random, then a male is chosen at random. whether or not the male “proposes” to the female is determined based on the relative (fork) lengths of each individual. If the female “accepts” the male’s proposal mating occurs and each of the females’ eggs receives one allele from each parent at each of the 20 loci that determine the offspring’s probability of maturing within a specific length-interval Once mating occurs, the female is removed from the class of potential mates and placed in the spawner population. Males remain in the candidate pool and may mate with one or more other females. The equations for fecundity and mate choice are described in Supplement File S1, Section A1 and parameter values are listed in Supplementary File S2, Tables H and I.

#### Egg development and parr emergence

All fertilized eggs are assumed to be the same size and survive to become parr (post-emergent fry) on May 1 according to a random binomial survival probability, segg. Surviving parr are assigned a common fork length of 37 mm and weight of 0.3545 grams. The value of segg is listed in Supplementary File S2, Table J.

#### Parr-to-smolt growth and survival

Parr survive to become smolts on July 1 according to a Ricker density-dependent function and grow according to the growth component of the McGurk equation, as described in the following section and Supplementary File S1, A5. Parameter values are given in Supplementary File S2, Tables K and L.

#### Post-smolt growth and survival

Smolts and sub-adults up to the age/length of maturity grow and survive according to the McGurk integrated growth-and-survival model [45]. The equations of the McGurk model are derived from the allometries of growth and mortality. The growth coefficient is the value of GR assigned on the basis of the individual’s maturation genotype (S^φ^) that determines which bivariate normal distribution of growth rate (GR) and target maturation length (LMat) the individual inherits (described in Supplement Files S1, **Logic and governing equations for genetic assignment of growth and length)**. Details of the McGurk model are provided in Supplementary File S1, A5, and parameter values are listed in Supplementary File S2, Table L.

Daily growth rates and maturation lengths are assigned such that individuals genetically programmed to mature at smaller sizes and younger ages have higher growth rates than individuals programmed to mature at longer lengths (and greater weights) and generally older ages. This is expected for most semelparous anadromous salmon [24, 57]. Analysis of 1000 simulated values of each of the five GR/WMat BVNs demonstrate that the parameterization of these distributions produce the expected negative correlation between maturation age and length on the one hand and daily growth rate on the other (Table 2).

**Table 2.** Correlation coefficients between maturation length-at-age and daily growth rate of each mature age class and of the entire set of age classes combined from 1000 simulated values of GR and LMat for each age-/length-at-maturation class (L2 to L6).

| Age-/length class: | Corr(Lmat, GR): |
| --- | --- |
| All | -0.901 |
| L2 | 0.899 |
| L3 | 0.902 |
| L4 | 0.894 |
| L5 | 0.906 |
| L6 | 0.907 |

The McGurk integrated growth and mortality model generates stage-to-stage survival rates that incorporate the effects of both size and rate of growth between life-history stages. The integration of the two effects quantifies the trade-off between 1) reducing future predation risk by rapid growth in the near term in order to obtain a larger size and 2) increasing the mortality risk in the short run due to faster growth that increases predation risk and possible metabolic risks due to reduced tissue maintenance. All else being equal, individuals of a given size that grow relatively fast incur greater mortality rates than same size individuals that grow slower; and, conversely, individuals of small size foraging at a given rate incur greater mortality rates than larger individuals that forage at the same rate in the same or similar environment.

#### Harvest, maturation and spawning

Harvest is modeled to occur annually immediately prior to spawning November 1. Harvest of each vulnerable individual is modeled randomly based on a global exploitation rate (hg) that applies equally to all vulnerable individuals. The global exploitation rate may be modified for each of 5 contiguous length intervals of mature individuals and each of 4 contiguous length intervals of immature individuals. The intervals may be chosen to correspond to the range of lengths of the majority of each age at maturity (2 to 6) and each age of immature individuals (2 to 5), or be otherwise chosen for length intervals for which different harvest rates are applied. For each interval, the target harvest rate is determined by multiplying the global exploitation rate by a user-chosen length-interval state variable *v* between 0 and 1 to yield the actual harvest rate ha (= hg*v) to which individuals in the specific length-intervals are vulnerable. This provides the flexibility to model harvest scenarios with different length interval and maturation status-specific harvest vulnerabilities, which in turn allows modeling of different harvest management regimes and gear selectivity. Specifically, terminal fishery scenarios are modeled only on mature adults, with harvest vulnerabilities of immatures of all length intervals set equal to 0. Mixed-maturation fishery scenarios can explore a range of vulnerabilities of both mature and immature individuals, including minimum length thresholds and drop-off and non-retention mortalities of sub-legal size matures and immatures.

Mature individuals that survive harvest move to the mating/spawning population at the end of each annual time-step. Unharvested immatures undergo growth and survival to the next time step.

### Model initialization

#### The unfished equilibrium population

The model is initialized with a mix of mature individuals of various ages and sexes in their target equilibrium proportions, numbering 1000 in total. For both sexes, the mean lengths of each age are shown in Table 1. These initial abundances and the proportions of both sexes combined are listed in Supplementary File S2 (B.6, Tables O, P, and Q). Individuals in each age-/length-class are initialized with distinct age-/length-specific mean alleles frequencies at the 20 loci controlling the maturation probabilities. Supplementary File S3 provides additional information.

#### Heritability of maturation age and length

Heritabilities for life-history traits are known to be smaller than heritabilities for morphological and behavioral traits [16, 58, 59]. This is consistent with the expectation that selection should deplete standing genetic variation in fitness [60, 61]. It is also expected from the bet-hedging parameterization of the genetic control of daily growth rates and size at maturity. We estimated heritabilities of smolts and recruited adult spawners of the same cohort for length– and age-at-maturation, and daily growth rate (GR). Estimates were made separately for each sex and for both sexes combined using mid-parent/offspring regression. Heritabilities were reasonably low, ranging from 0.07 to 0.20 (Supplementary File S2, Table R).

### Harvest scenarios

The model provides for the ability to model harvest with either a minimum threshold length limit, as appropriate to a troll fishery, or a fishery with both a lower and an upper size limit, as may occur in a gillnet fishery. To provide a robust evaluation of the impact to Chinook demographics from both troll and gillnet harvest in terminal and mixed-maturation fisheries, we evaluate 6 harvest scenarios that explore the main features of these 2 kinds of harvest regimes. All scenarios were run with the same seed for the random number generator and initiated with 100 years of no harvest (pre-fishery scenario), to attain the adult equilibrium spawner abundance and age/sex composition. This was followed by 25 years of harvest with a constant global harvest rate, hg, and a set of vulnerabilities to the global rate, v, of matures and immatures in each maturation length interval ranging from *v* = 0 for completely invulnerable to v = 1.0 for fully vulnerable. This yielded one or more length-interval and maturation-state-specific harvest rates ha (=hg*v). Catch of individuals vulnerable to harvest was modeled in all scenarios by drawing a uniform random number, u, on [0, 1] and considering an individual caught if *u* is less than or equal the value of ha to which the individual is vulnerable.

The 6 scenarios consist of 3 basic troll fishery scenarios (H1, H3, and H5) and 3 similar basic gillnet fishery scenarios (H2, H4, and H6). The 3 types of fisheries modeled for both troll and gillnet scenarios are, in order: 1) a terminal fishery harvesting matures at a global harvest rate (hg) that achieves maximum sustained yield in the total weight of the catch (scenarios H1 and H2); 2) a mixed maturation fishery harvesting immature and mature Chinook at a global harvest rate (hg) that achieves maximum sustained yield in the total weight of the catch (scenarios H3 and H4); and, 3) a terminal fishery harvesting matures at a global harvest rate that approximates the total weight of the MSY catch of the corresponding MM scenario (scenarios H5 and H6). We refer to this latter harvest rate and corresponding harvest scenario as the “mixed-maturation-MSY-catch-weight-equivalent (MMCWE)”. For scenarios H1 to H4, the approximate global MSY harvest rate (hg) was estimated by trial-and-error by running each 150-year simulation with the global harvest rate (applied in years 101 to 125) set to several different values over intervals of 0.05 (e.g., 0.60, 0.65, 0.70) and choosing as the global MSY harvest rate (Hmsy) the value of hg that generated the largest total weight of the catch. In the case of scenarios H5 (troll MMCWE) and H6 (gillnet MMCWE), the global harvest rate was further refined at intervals less than 0.05 as necessary to achieve a close approximation to the corresponding MM scenario total catch weight.

Scenarios H1 to H6 were first run with no post-smolt environmental variation, simulating the average environmental conditions affecting density-independent growth and survival. We then ran each scenario with environmental variation affecting daily growth rates and annual survival rates. Details are described in Supplementary File S1, A7 and A8. Parameter values are described in Supplementary File S2, B5.

The features of each of the 6 scenarios are listed in Table 3. For terminal fishery scenarios, vulnerabilities (v) of all immatures were set equal to 0, and all matures were set to 1.0. For mixed-maturation fishery scenarios, v of all matures and immatures were set equal to 1.0. We provide further details in the Discussion.

**Table 3.** Harvest scenarios H1 through H6 with their global harvest rates and vulnerabilities of immature fish. hg is the global harvest rate; v is the length range-specific modifier of the global harvest rate applied to immature and mature fish. The actual harvest rate for individuals in length ranges for which v is less than 1.0, ha, is equal to hg*v.

| Scenario | Type | Gear<br>Type | $hg$ | Immature Vulnerability<br>($v$) |
| --- | --- | --- | --- | --- |
| H1 | Terminal MSY | Troll | 0.65 | 0.0 |
| H2 | Terminal MSY | Gillnet | 0.70 | 0.0 |
| H3 | Mixed-maturation<br>MSY | Troll | 0.45 | 1.0 |
| H4 | Mixed-maturation MSY | Gillnet | 0.50 | 1.0 |
| H5 | Terminal MMCWE | Troll | 0.42<br>5 | 0.0 |
| H6 | Terminal MMCWE | Gillnet | 0.44 | 0.0 |

Global harvest rates (hg) in scenarios 1, 2, 3, and 4 are estimates of the MSY harvest rate. Global harvest rates in scenarios 5 and 6 are estimates of the harvest rate in terminal fisheries that produce approximately the same total catch weight (mixed-maturation-MSY-catch-weight-equivalent/ MMCWE) as the corresponding mixed-stock MSY fishery (scenarios H3 and H4, respectively).

Since we model harvest as occurring immediately before unharvested fish spawn, annual natural mortality during the final year of life of mature fish implicitly includes natural mortality incurred during spawning migration that occurs after passing the fishery. We consider this to be a reasonable simplification and discuss it further in the Discussion.

#### Fishery scenario H1. Terminal MSY troll fishery harvest on mature adults

Harvest scenario H1 simulates a terminal fishery harvesting mature adults equal to or greater than 610 mm (∼24 inches) fork length at a global harvest rate (hg) that achieves maximum sustainable yield (MSY) in total weight of the catch. This excluded harvest of most age 2 mature males (mean fork length 450 mm (Table 1, Fig 2) plus 3 standard deviations for CV length = 0.09*mean: 572 mm. This established a baseline against which to compare MM fishery scenarios involving harvest of mature and immature Chinook.

We chose 24 inches (610 mm) as the minimum fork length, as this is the size limit for the tribal treaty troll fishery off the northwest Washington Coast [63]. We feel it provides a reasonable, and likely conservative, estimate of the minimum fork length of Chinook vulnerable to capture by troll gear in the early years of mixed stock coastal troll fisheries to which our model population would have been vulnerable.

#### Fishery scenario H2. Terminal MSY gillnet fishery harvest on mature adults

Harvest scenario H2 is the gillnet counterpart to scenario H1. H2 simulates a terminal fishery harvesting mature adults at the global harvest rate (hg) within the length interval [783 mm, 1158 mm], and at ha = 0.1* hg at lengths below 783 mm and ha = 0.4* hg at lengths above 1158 mm. The MSY harvest rate is estimated by trial and error by simulating the harvest scenario at several different values of hg at intervals of 0.05, starting at 0.60 as for scenario H1. For further details, see the section **parameterization of gillnet scenarios** below and Supplementary Files S1, A9, and S2, B8.

#### Fishery scenario H3. Mixed maturation MSY troll fishery

Harvest scenario H3 simulates a mixed maturation troll fishery on mature and immature Chinook with a minimum fork length of 610 mm (vulnerability of age 3 and older = 1.0). Individuals aged 3 to 6 of both sexes greater than 610 mm FL were fully vulnerable to the gear regardless of maturation state. This would result in only a small proportion of age 3 immatures and matures not being vulnerable to the fishery, essentially simulating a troll fishery with a minimum size limit of 610 mm FL. We also assumed that the 610 mm cut-off was perfect (knife-edge), in the sense that there was no by-catch of sub-legal fish. Thus, in effect, we model fisheries with gears that are perfectly selective for fish equal to or greater than the minimum size limit. The global harvest rate was set by trial and error to determine the rate that achieved the MSY harvest when age 3 to 5 immatures >= 610 mm FL are equally as vulnerable to the gear as mature (age 3 to 6) individuals.

#### Fishery scenario H4. Mixed maturation MSY gillnet fishery

Similar to scenario H3, this scenario searched for a value of hg that achieved the greatest total catch weight in a gillnet fishery in which both mature and immature Chinook were vulnerable to a gillnet as parameterized in scenario H2.

#### Fishery scenario H5. Terminal mixed maturation catch weight equivalent (MMCWE) troll fishery

This scenario searched for a global harvest rate (hg) that achieved a total catch weight approximately equal to that of the mixed maturation MSY troll fishery (H3), but in a terminal fishery in which only matures are harvested. This consideration provides the harvest rate that would be required of a terminal fishery replacing a MM fishery, but achieving the same average annual total catch weight. This scenario is identical to scenario H1 except that the global harvest rate was set by trial and error to achieve approximately the same total annual catch weight as scenario H3, (“catch-equivalent MSY harvest rate”).

#### Fishery scenario H6. Terminal mixed-maturation – catch weight equivalent (MMCWE) gillnet fishery

Similar to scenario H5 but for a terminal gillnet fishery, this scenario searched for a global harvest rate (hg) that closely approximated the total MSY catch weight achieved in the MM gillnet scenario H4.

#### Parameterization of gillnet fishery scenarios

Gillnets generally harvest individuals with fork lengths between a lower and upper size limit, at or near the global harvest rate. Individuals with lengths below the lower and above the upper limits are harvested at rates below the global rate due to lower vulnerabilities to the gillnet mesh (lower selectivities). In addition, gillnets generally impose different selectivities on lengths within the interval of greatest vulnerability, with the greatest selectivity (catchability) on intermediate lengths. Catchability generally drops abruptly below the lower length limit and more gradually above the upper limit. Below the lower length threshold catchability declines abruptly as smaller fish can swim through the mesh and are entangled less frequently than fish above the upper threshold. Above the upper length threshold, catchability gradually levels off above zero, because larger fish can still be caught from entanglement of the fins or operculum. Supplementary File S1, A9, Fig B shows a gillnet selectivity curve for the standard 8-inch mesh commonly used for Chinook. This curve was based on a Pearson function applied to an extensive data set for Yukon River Chinook salmon analyzed by [62].

We modeled gillnet catchability with an approach that is a compromise between using an explicit function like Pearson that assumes a selectivity of 0.0, and the lower and upper length threshold within which selectivity is 1.0 (e.g., [34]). Based upon the selectivity curve of our parameterization of the Pearson function of [62], we chose the interval [783, 1158 mm FL] as the length interval in which vulnerability was equal to 1.0. Selectivity on lengths below 783 FL was set to 0.10, and selectivity on lengths greater than 1158 were set to 0.40. (See Supplementary Files S1, A9 and S2, B8 for further details).

## Results

We describe and compare the results of the troll and gillnet harvest scenarios under no environmental variation for each of the three types of harvest scenarios.

To consider environmental variation, we repeated the above harvest scenarios (H1 to H6) adding bivariate normal random variation to the daily growth (GR) and annual survival (S) rates of smolts and post-smolts age 1 to 5 (as described in Supplement File S1, A8). None of these additional harvest scenarios produced results that differed substantially from the 6 primary scenarios in regard to the magnitude or direction of the differences between mixed maturation and terminal harvests. Consequently, we do not report them.

We provide graphical summaries that highlight specific impacts of each scenario on population life history, and summarize the results of each harvest scenario. Data tables of the primary results are provided in Supplementary File S4 and S5 Table.

### Comparisons of mixed maturation to terminal harvest scenarios

There are clear differences between the terminal and MM scenarios for both gear types in regard to the average weight of the catch, the age composition of the returning adult and spawner populations, (Figs 3 and 4) and in the mean age composition of the catch and spawner populations (Figs 5 and 6). In each case, MM fisheries shift the age composition of both sexes toward younger ages and smaller average catch weights (Fig 3).

**Fig 3.**
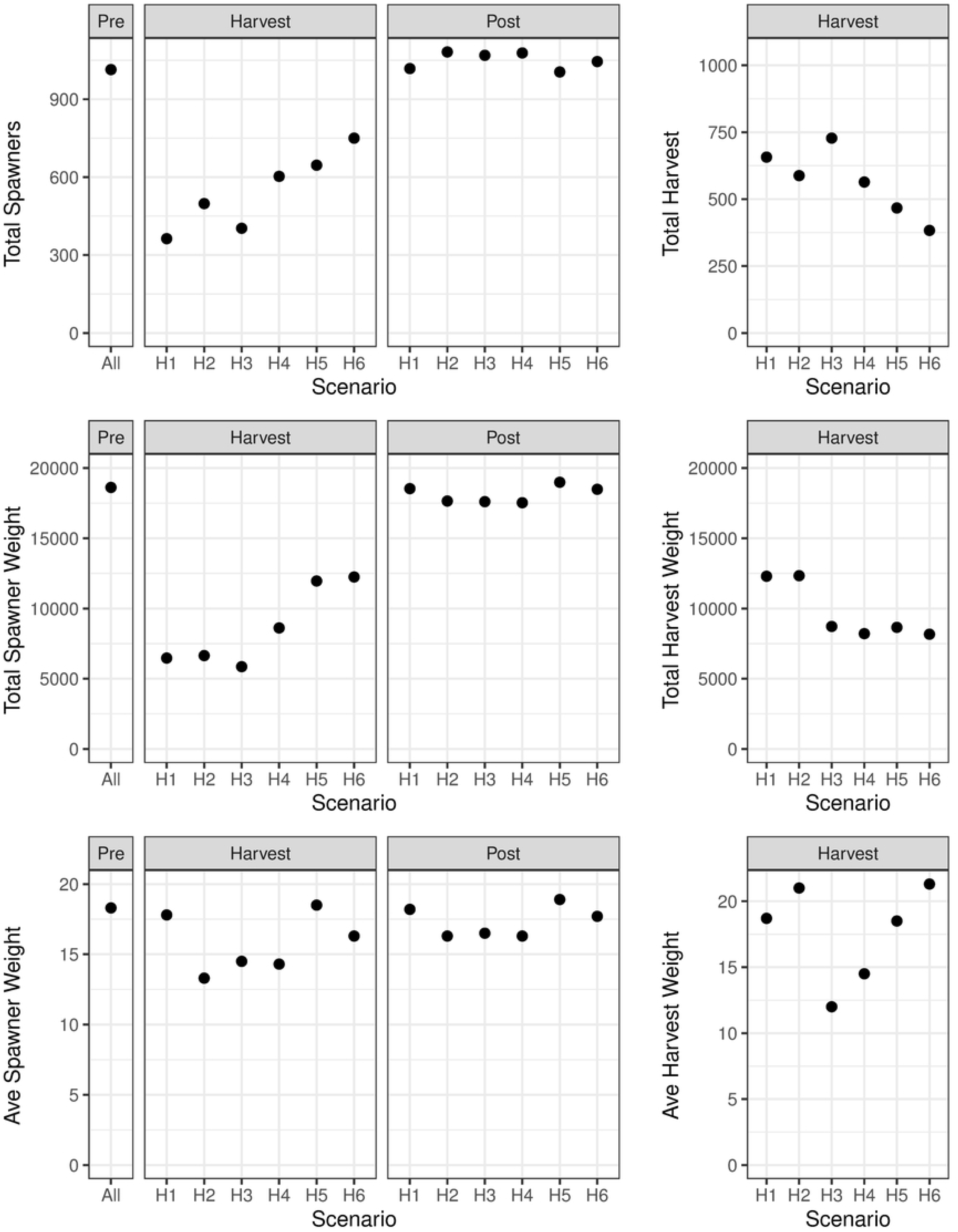
Mean number of total spawner and harvested numbers and weights, and average weights, for all 6 harvest scenarios for the harvest and post-harvest periods (simulation years 101 – 125 and 126 – 150, respectively). X axis labels: H1: Terminal troll; H2: Terminal gillnet; H3: Mixed maturation troll; H4: Mixed maturation gillnet; H5: Terminal mixed maturation catch equivalent troll; H6: Terminal mixed maturation catch equivalent gillnet. We do not show standard errors as they are very low for all groups, ranging from a maximum of 0.068 for terminal harvest scenario H1 and a minimum of 0.002 for pre-terminal spawner weight.

**Fig 4.**
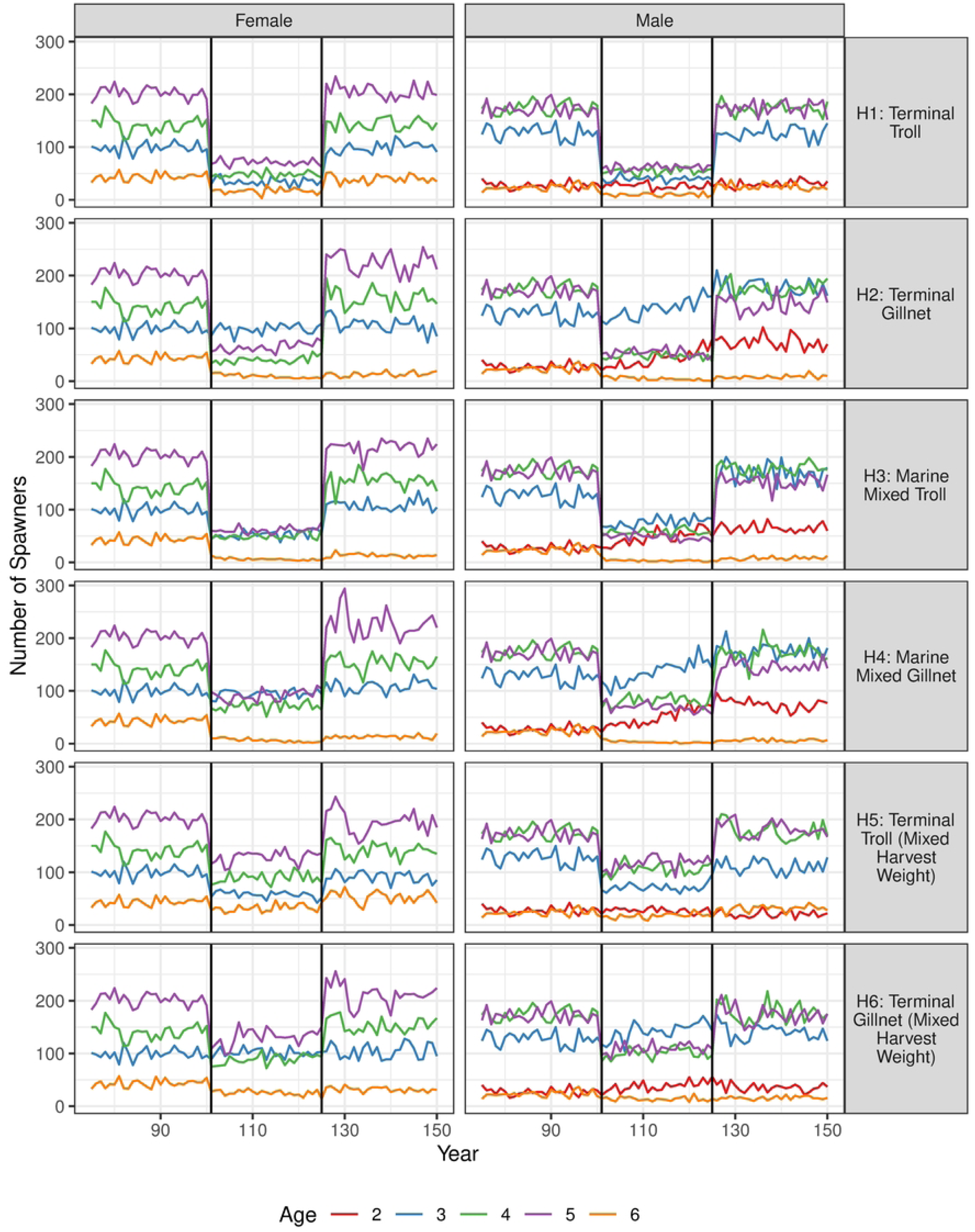
Time series of number of spawners by age and sex for simulation years 75 to 150 for all 6 harvest scenarios. Harvest period shaded in grey.

**Fig 5.**
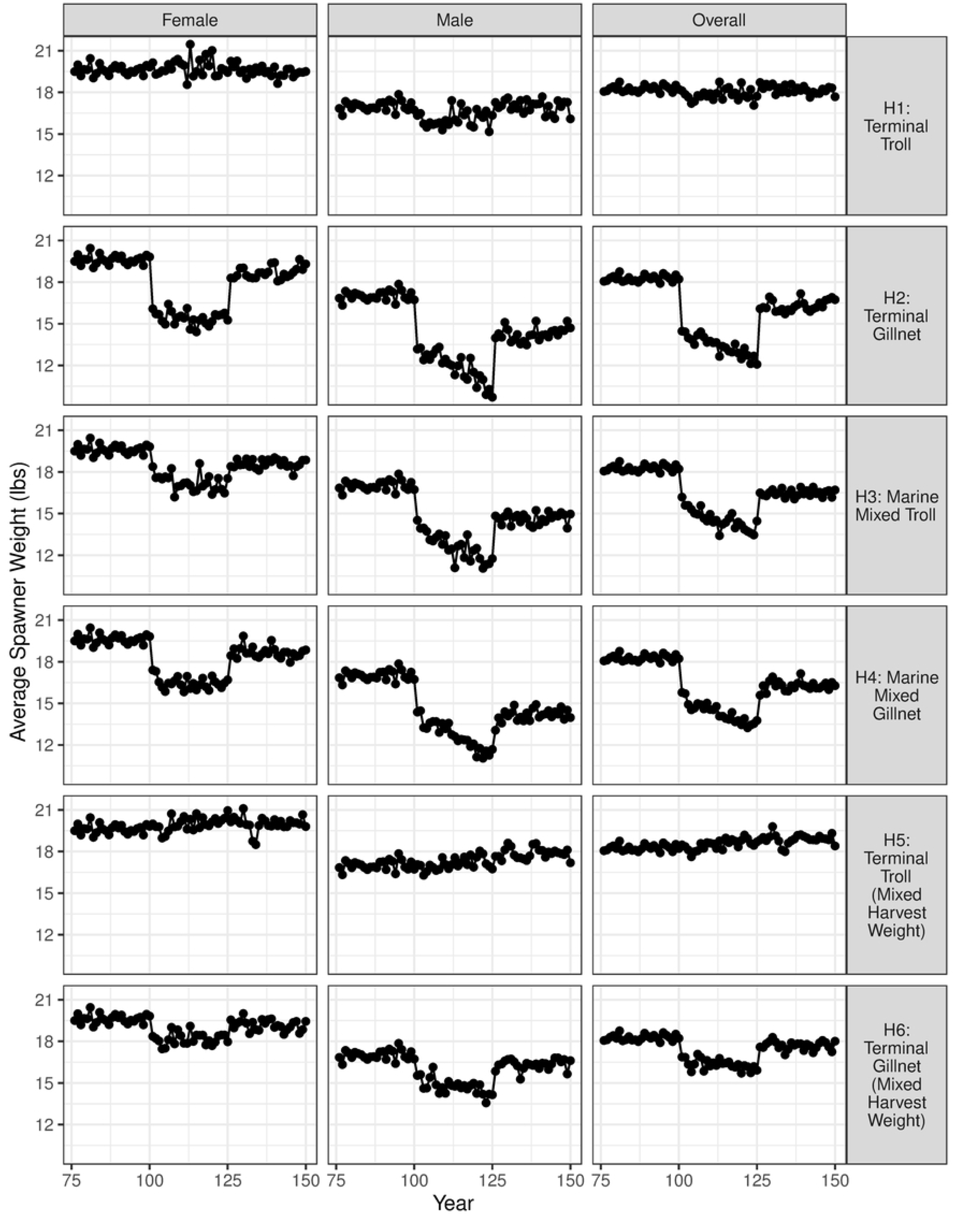
Average spawner weights for each harvest scenario. Harvest period shaded in grey.

**Fig 6.**
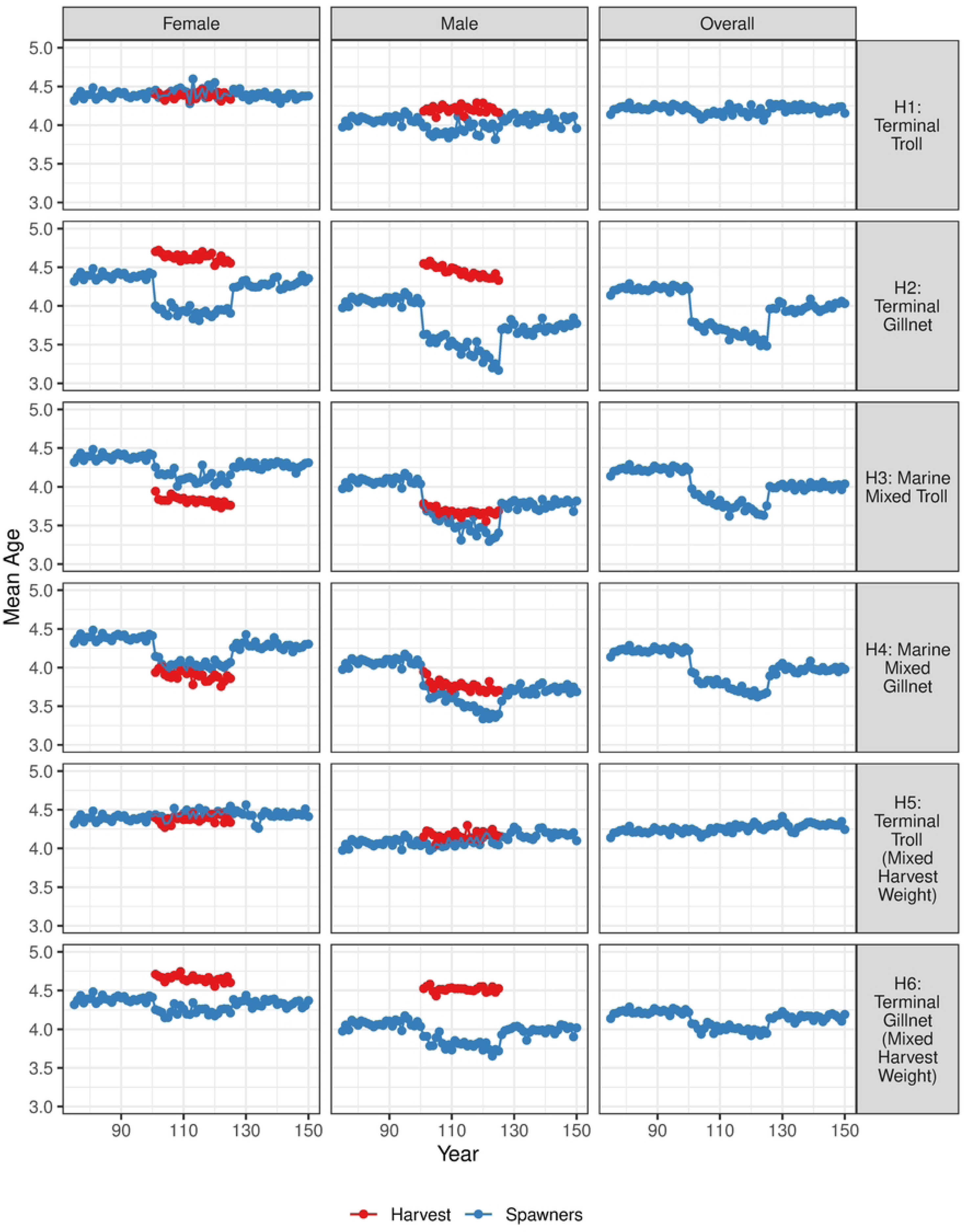
Average age of spawners and harvested Chinook for each sex for simulation years 75 to 150 for all six harvest scenarios.

In addition, the total weight of the catch at MSY in MM fisheries is 30% to 35% lower than in terminal MSY fisheries for both gear types (Fig 3). MM fisheries of both gear types significantly decrease the proportions of the 2 oldest age classes relative to the terminal fishery (Figs 4 and 7; Supplement File S4, Table C). MM fisheries also decrease the average weight of the spawning population (Fig 5), and thereby impair the ability of the harvested population to recover in the direction of the pre-fishery age composition compared to the terminal scenarios (Figs 4 and 5; Supplement Files S4, Table D). This is primarily due to the legacy of the harvest of immature individuals in the MM fisheries.

**Fig 7.**
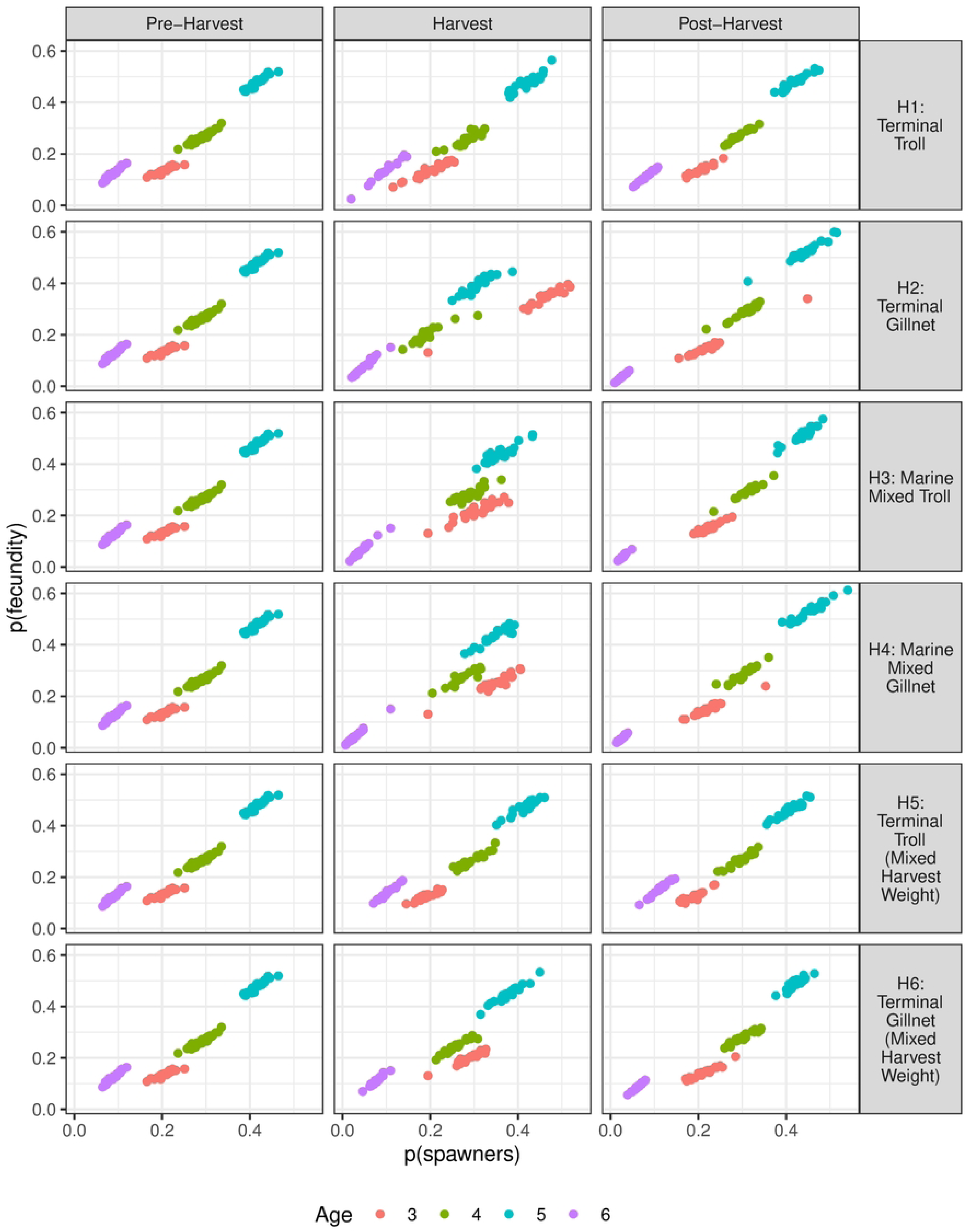
Age-specific proportions of total egg deposition vs. age proportions of spawners by harvest scenario.

Importantly, over the 25-year harvest period the 2 terminal MSY scenarios (H1 and H2) achieve considerably greater total and average harvest weights than the MM MSY scenarios (Figs 3 and 8; Supplement File S4, Table A).

**Fig 8.**
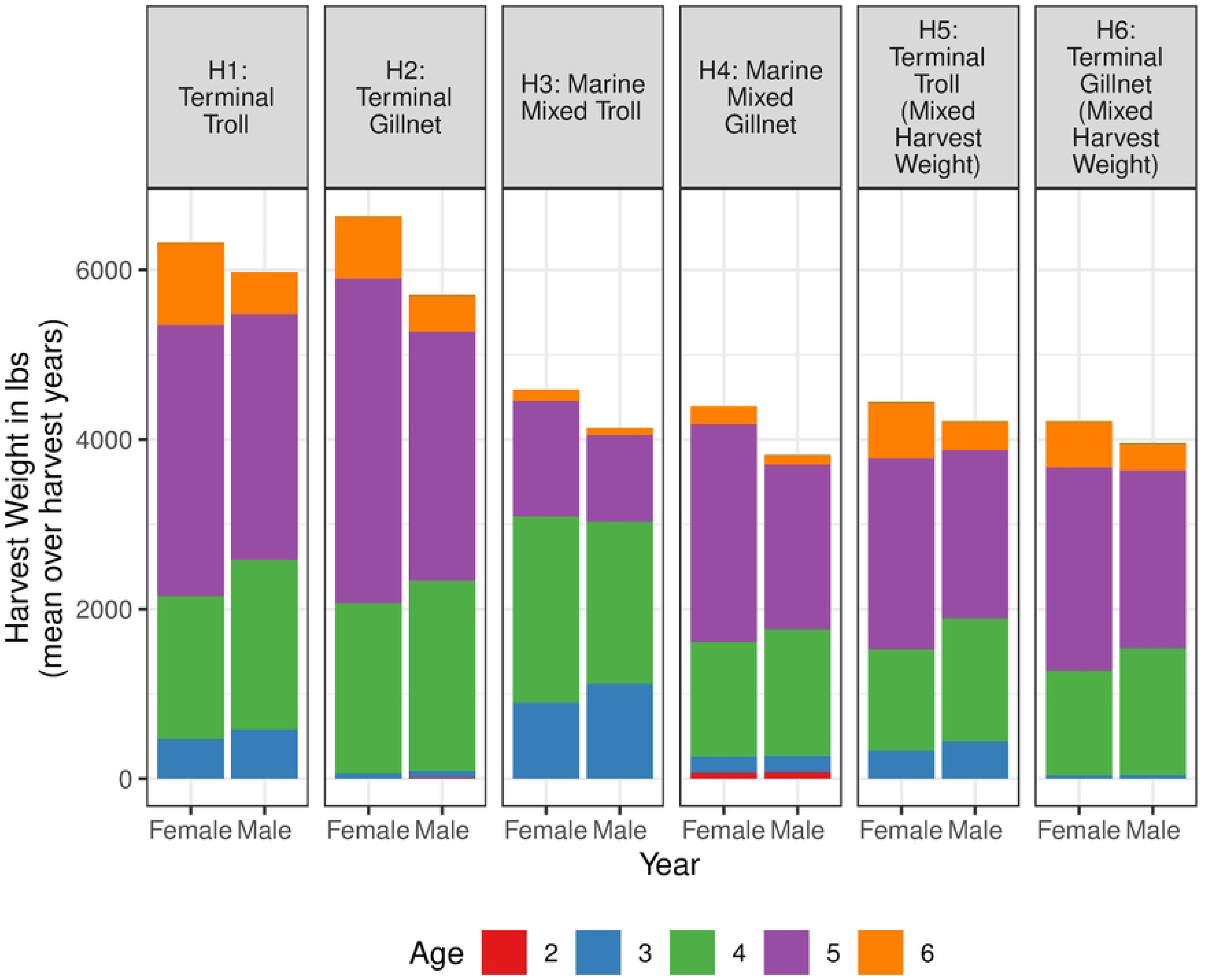
Total weight of harvest for all 25 years for each sex in all six harvest scenarios.

With respect to the sustainability of the fishery regimes, the terminal catch-equivalent MM fisheries (H5 and H6) achieve the target total catch weights of the MM MSY scenarios with fewer and larger fish than the corresponding MM MSY fisheries. Additionally, they do so with a spawning population that has a larger average size, weight, and age than the mixed-stock fisheries (Figs 3, 5 and 6).

### Troll scenarios versus gillnet scenarios

There are biologically meaningful differences in average spawner and catch weights between the 2 gear types in each of the three types of fishery scenarios. In the terminal MSY troll fishery scenarios (H1), the average weight of spawners during the harvest period is one-half lb smaller than the pre-harvest average (17.8 versus 18.3). In the terminal MSY gillnet scenario (H2), the average weight of spawners during the harvest period is a full five lb smaller than the pre-harvest average (13.3 versus 18.3; Figs 3 and 5). Comparing between fisheries, the average size of harvested fish in the terminal MSY gillnet fishery is more than 2 lb greater than in the corresponding troll fishery (21.0 lb versus 18.7 lb; Figs 3 and 5).

In the mixed-maturation MSY scenarios (H3 and H4), both total and average catch weights of each gear type are significantly below their terminal fishery counterparts (Fig 3). However, there are some interesting differences between the 2 gear types. In the mixed-maturation troll scenario, the total number of fish caught is greater than in the gillnet scenario (728 versus 564; Figs 3 and 9), but the average weight of the catch is smaller (12.0 versus 14.5 lb; Fig 3). This is due to the greater selectivity of the gillnet for larger, older fish than the troll scenario, which captures more younger-age, immature fish. Correspondingly, there are more spawners in the gillnet scenario (603 versus 403), but average spawner weights are similar (14.5 versus 14.3, respectively; Fig 3). This is due to the troll fishery catching more fish across the size range, including more immatures, than the gillnet (432 weighing an average of 9.5 lb versus 226 weighing an average of 5.3 lb, respectively (Fig 9)). This leaves fewer spawners on average than the gillnet fishery (Fig 4). In both cases, the average spawner weights are 4 lb smaller than the pre-harvest average.

**Fig 9.**
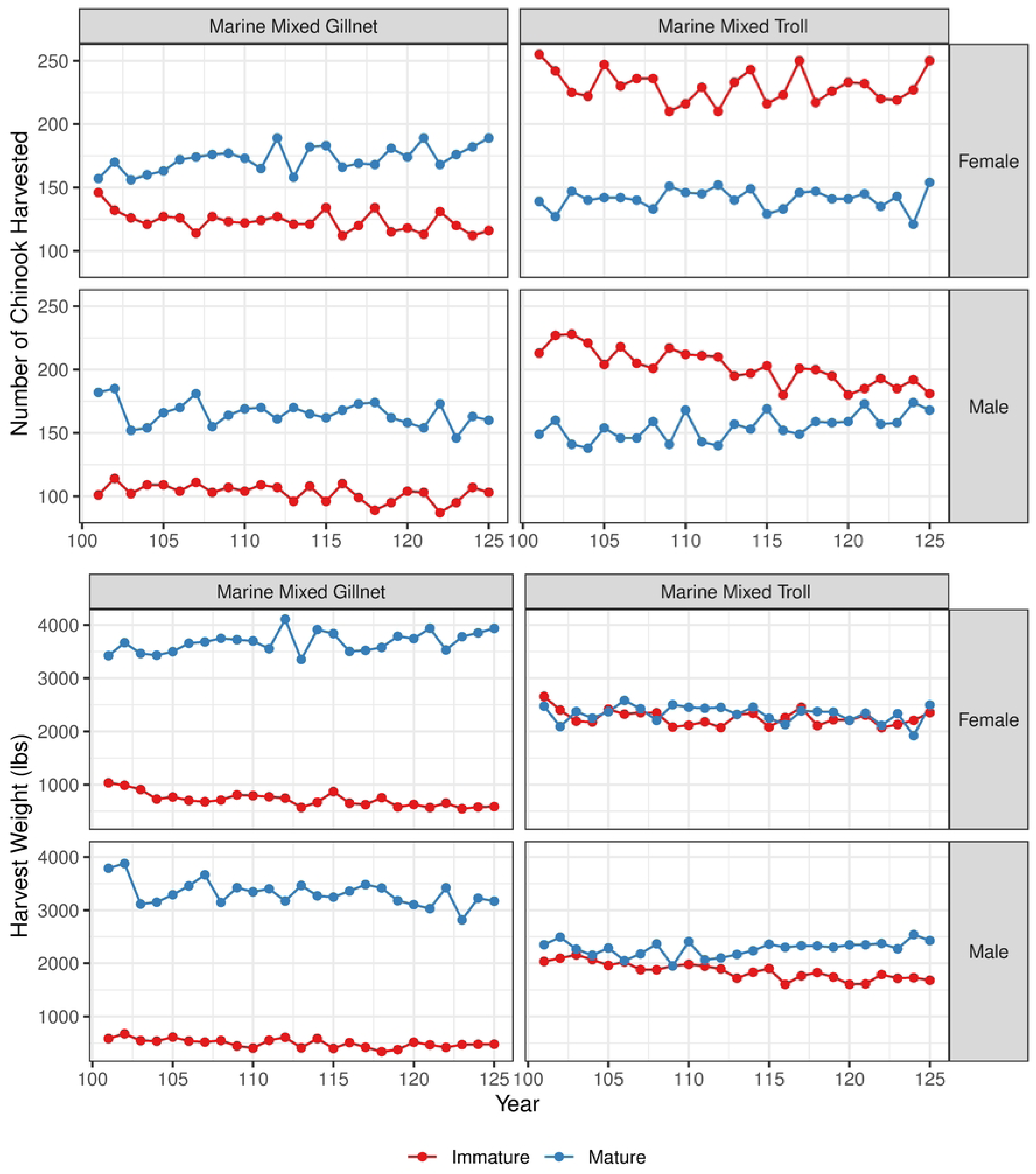
Total number and weight of Chinook of each sex harvested in the mixed maturation troll and gillnet fishery scenarios.

In the MMCWE fishery scenarios (H5 and H6), the terminal troll scenario achieves an average spawner weight of 18.5 lb, compared to the pre-harvest average of 18.3, and an average catch weight of 18.5 lb, which is similar to the average catch weight of the terminal MSY troll scenario. Total number of spawners is nearly 300 more than in the terminal MSY fishery and more than 200 than the MM troll scenario. The gillnet scenario achieves an average weight of spawners of 16.3 lb, and an average catch weight of 21.3 lb, and 100 more total spawners (750) than the troll scenario (Fig 3; Supplement File S4, Table A).

The mean age of spawners in the MMCWE troll scenario is slightly, though not significantly, greater than in the terminal MSY troll scenario and the mean spawner weights in each period are equal to or slightly greater than the pre-harvest means (Fig 3). In the MMCWE gillnet scenario, the mean ages in the harvest and post-harvest periods are lower, but close to the values in the terminal MSY troll scenario (Fig 6), while the mean weight of spawners in these two periods are 1 to 2 lb smaller than in the two terminal troll scenarios (Fig 3; Supplement File S4, Table D).

### Harvest impacts to egg deposition

Harvest impacts are also reflected in the total and age-specific egg deposition and in the proportional contribution of female spawners of each age. For all 6 harvest scenarios, we provide summary data for the proportion of total egg deposition contributed by female spawners of each age (3 to 6) relative to the proportion of each female age in the total female spawner population. Fig 10 shows the total egg deposition for simulation years 75 to 150 for all six harvest simulation scenarios. data is provided in tabular form in Supplementary File S4, Tables E and F.

**Fig 10.**
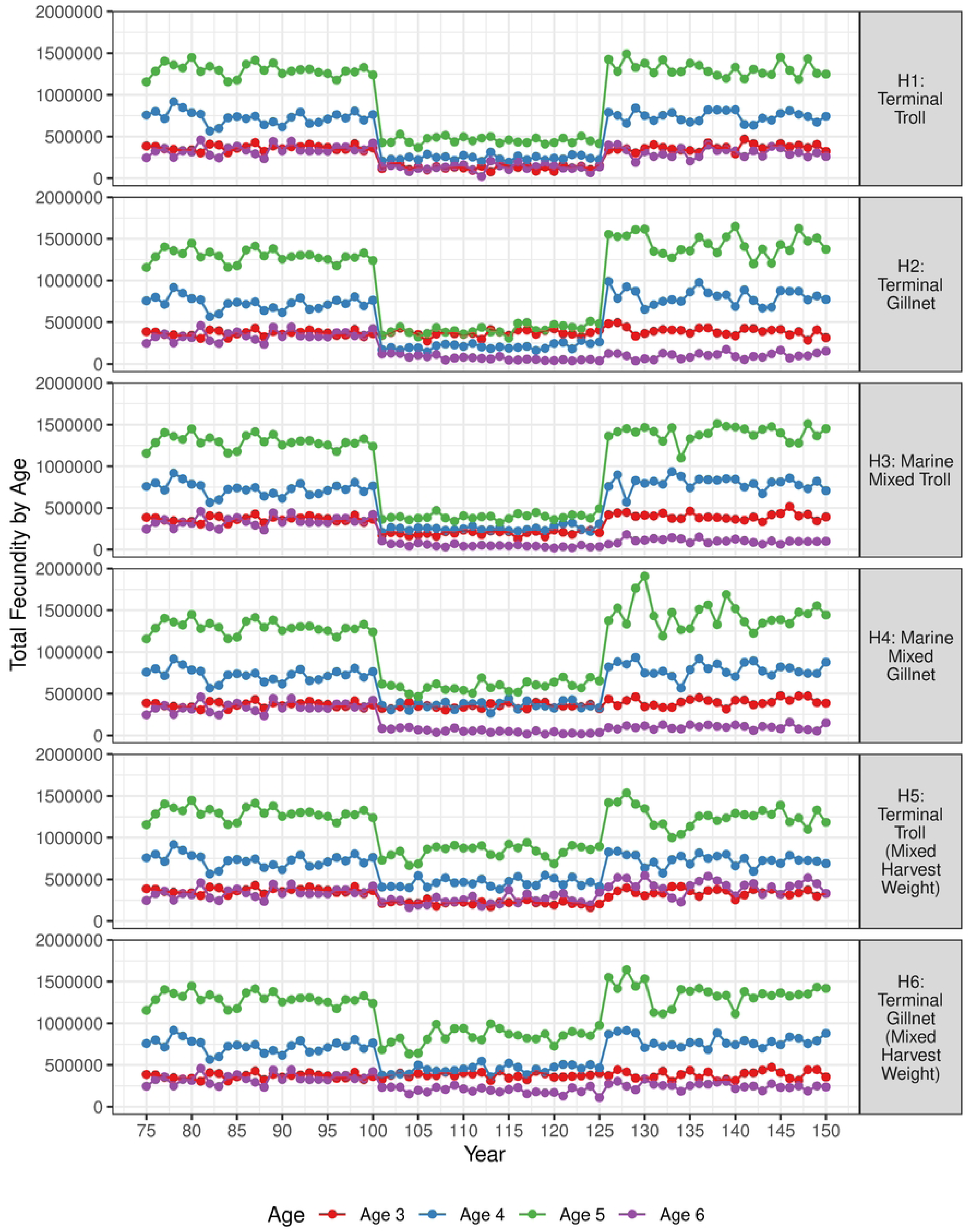
Age-specific total spawner egg deposition by harvest scenario for simulation years 75 to 150.

The reference terminal harvest scenarios (H1 and H2) reveal clear differences in the impact on the age structure of the spawning population between the troll and gillnet. The age composition in the terminal troll scenario is noticeably less skewed toward younger ages (3 and 4) relative to the pre-harvest period, and compared to the terminal gillnet scenario (Fig 10; Supplement File S4, Table C).

An illustrative comparison of the impact of each scenario on the structure of the spawning population is the proportion of age 3-to-6-year-old spawners and the proportion of total egg deposition contributed by each. In the gillnet scenario (H2), the proportion of age 3 female spawners increases and the proportion of age 4, 5, and 6 female spawners decreases relative to the pre-harvest period. This change is mirrored by the proportion of total egg deposition contributed by age 4 and 5 spawners, but less so in the case of age 6 spawners (Fig 10, middle panel; Supplement File S4, Table C). During the pre-harvest period, age three females comprise 15% to 25% of the spawning population but contribute less than 18 percent of the total egg deposition. During the harvest period, the proportion of age three females in the spawning population rises above 40% and contribute 30% to 40% of the total egg deposition. Both numerically and in terms of contribution to total egg deposition, the gillnet harvest renders the population during the harvest period more dependent on age 3 spawners, which likely have lower overall fitness measured as recruits-per-spawner than the 3 older ages. During the post-harvest recovery period, age 5 spawners increase significantly in both proportions relative to the pre-harvest period, age 3 and 4 spawners recover to near their pre-harvest levels, but age 6 spawners fall below pre-harvest levels and thus fail to rebuild during the 25-year period.

In contrast, there is little significant change in either proportion of the 4 age classes in the terminal troll scenario (H1). Thus, harvest across the entire size range (fork length) of mature adults vulnerable to the troll gear has little impact on either the proportion of each age class in the total spawning population, or in the proportional contribution to egg deposition relative to the pre-harvest period. Consequently, there is little disruption of spawning age structure, and less correction during the 25-year recovery period.

In the mixed maturation fishery scenarios, the situations are similar between the 2 gear types (H3 and H4) (Fig 10; Supplement File S4, Table C). Both scenarios result in a marked reduction in the proportion of age 5 and 6 female spawners and corresponding increases in the proportion of ages 3 and 4 female spawners, relative to the pre-harvest proportions. Pre-harvest, ages 5 and 6 combined make up 51% of total spawners and ages 3 and 4 combined 49%. In the mixed maturation troll harvest period, ages 5 and 6 combined represent 40% and ages 3 and 4 60% of total spawners. In the mixed maturation gillnet harvest period, ages 5 and 6 represent 37% and ages 3 and 4 63% of total spawners (Fig 10; Supplement File S4, Table C). Both gear types effect similar reductions in the proportional contribution to total egg deposition of age 6 spawners, similar increases in the proportional contribution of age 3 spawners, and little alternation in the proportional contributions of ages 4 and 5 spawners (Fig 10). There is little difference between the 2 scenarios during the post-harvest recovery period. In both scenarios, age 6 females lag behind the pre-harvest proportions. The 3 younger age classes recover to near pre-harvest period levels, though there are some slight differences between the 2 gear types.

Age composition of the spawning population in the 2 terminal catch weight equivalent harvest scenarios (H5 and H6) shows considerably less alternation from the pre-harvest age structure, than the corresponding mixed-maturation scenarios. However, the MMCWE gillnet scenario produces an increase in the proportion of age 3 spawners that does not occur in the corresponding troll scenario. The troll scenario (H5) retains the pre-harvest average proportions of ages 5 and 6 in the total spawning population (0.522 versus 0.518 The average proportions of age 5 and 6 spawners in the MMCWE gillnet scenario is 0.452 (Fig 10; Supplement File S4, Table C), an improvement over the average in the mixed maturation scenario (H4) of 0.372, but still below the pre –fishery average. However, both scenarios permit a recovery to at (troll) or near (gillnet) the pre-fishery spawner age proportions over the 25-year post-harvest period (Supplement File S4, Table C).

To provide further perspective for evaluating the impacts of the harvest scenarios on age-specific spawner contributions to population growth, we calculated the reproductive values of female spawners (ages 3 to 6) for a representative average deterministic population projection matrix. For interested readers further details are provided in Supplement File S1, section A6.

## Discussion

### Size-versus age-overfishing

Our results show that in mixed maturation fisheries (where mature and immature individuals attain lengths that render them equally vulnerable to the fishing gear), mortality in MSY harvests imposes biological effects that are likely to compromise a population’s resilience and adversely affect potential yield. These effects are primarily demographic in the short-term extending one to several generations, the period of time over which we conducted our simulations. Biological effects occur from shifting the population’s size structure toward smaller lengths at maturity (i.e., size-overfishing) and, consequently, a younger adult age structure (Figs 3, 4, 5, 6, 10; Supplement File S4, Table C). These shifts occur independent of and in addition to, any potential genetic effect due to selection on growth or maturation rates. The shift toward a younger age structure is, therefore, an indirect effect of size selection. Note that the shift to smaller sizes is not a result of selection on size-at-age, since our model does not allow the genotype-specific maturation probabilities to evolve in response to selection on age or size. Rather, the shift to smaller size is simply the result of selection on the fitness of growth rates and lengths-at-maturity that result from the size selection of the fishing regime; it is fundamentally a demographic effect.

Thus, our model results in this respect are conservative. We expect that if selection on the genotype-specific maturation probabilities (daily growth rate and maturation weight/length) were incorporated into our model (via no or a reduced degree of diversified bet-hedging), selection toward smaller sizes and hence younger ages, would increase the shift in the directions observed. For example, if heritabilities for growth rate and size-at-maturity were higher, such that age x parents had a very high (e.g, >0.75) probability of producing offspring with growth rates and size-at-maturity identical to their own, and if size-assortative mating were much stronger than our current parameterization, older-aged/larger-size adults would have a very low (perhaps zero) probability of producing offspring with higher growth rates and smaller size-/younger age-at-maturity. Younger/smaller-size adults would have similarly low probabilities of producing offspring with lower growth rate and larger size-older age-at-maturity. Under exploitation rates as high as (or likely even less than) those observed in our terminal and mixed maturation MSY scenarios, there would be a high probability of losing larger, older adults altogether from the population due to strong selection for faster growth rates and smaller size-/younger age-at-maturity.

By incorporating diversification bet-hedging in our parameterization of the genetics of maturation (a form of genetic and environmental canalization that prioritizes geometric mean fitness rather than arithmetic mean fitness [46–50]) our model population retains the ability to produce all growth rates and maturation sizes and ages of the pre-harvest population under a range of realistic exploitation rates. However, given our focus on time periods relevant to management, (25 years or 5 to 6 generations), we chose not to incorporate the added complexity of allowing selection to operate on growth rates and/or maturation lengths. In any case, our results strongly suggest that incorporating evolution of the genetics controlling maturation would not affect the generality of our results regarding the benefits of terminal fisheries and the detrimental impact of mixed-maturation fisheries.

The demographic impacts that occur under mixed maturation harvest also have suboptimal effects on the total numbers and weight of the catch (Fig 8; Supplementary File S4, Table A). In the MM troll scenario (H3) where immature and mature fish 610 mm FL and larger are equally vulnerable to the global harvest rate, immature Chinook comprise 59% of the total numbers caught, and 47% of the total weight of the catch (Fig 9). In the MM gillnet scenario where immature and mature fish are also equally vulnerable to the global harvest rate, but the gear is disproportionately selective for larger, older fish, immature Chinook comprise 40% of the total number caught and 15% of the total weight of the catch (Fig 9). Importantly, we did not include release mortality on sublegal catch or drop-off mortality on legal and sublegal size fish in the mixed maturation scenarios, (phenomena that are common in marine, particularly troll, fisheries), again emphasizing the conservative nature of our results.

As noted in the Results, the average weight of spawners in both mixed maturation scenarios are reduced by roughly 4 lb from the pre-harvest average (Figs 3 and 5), the proportions of the 2 oldest spawner ages are reduced by more than 10% (Fig 7; Supplement File S4, Table C), and the mean age of spawners is reduced by nearly half a year from the pre-harvest average (Fig 6).

### Demographic effects of the two gear types on population structure

We found large differences in the effects of harvest on the size and age structure of the adult population between the 2 gear types in both mixed maturation and terminal fishery scenarios. These were due to the cutoff lengths for the selectivities of gillnet fisheries that we chose on the basis of Bromaghin’s analysis of the selectivity of the 8-inch mesh typically used for Chinook.

In troll scenarios, individuals at or above the minimum size threshold tend to be harvested close to their proportions in the total population vulnerable to harvest. Gillnets, by contrast, harvest larger, older individuals in proportions greater than their proportions in the total vulnerable population. Thus, individuals in ages 4 to 6 were harvested to a greater extent than ages 2 and 3, which were only harvested incidentally. This was due to two features of our parameterization.

First, our upper limit of 1158 mm FL was close to the largest length of the oldest (age 6) length class, so there were few individuals subject to the lower selectivity on lengths greater than 1158. Second, our lower limit of 783 mm meant that all age 2 individuals and most age 3 individuals were subject to the lower harvest rate (Supplementary file S1, Fig B). Thus, fish smaller than this lower limit consisted mostly of ages 2 and 3.

A related feature of the terminal troll harvest scenarios (H1 and H5) in comparison to the 2 mixed maturation (H3 and H4) and the 2 terminal gillnet scenarios (H2 and H6), is a negligible alteration of the mean age of the spawning population relative to the mean age at the unfished stochastic equilibrium (Fig 3; Supplement File S4, Table D). This is due to 2 features of these scenarios: first, the minimum length limit (610 mm) that we chose for the troll scenarios results in over 95% of all mature females and males age 3 and older being fully vulnerable to the fishing gear. Second, all vulnerable individuals are equally and randomly vulnerable to the global harvest rate and are therefore harvested (on average) in direct proportion to their age/length-interval-specific abundance, which tends to preserve the initial proportions between ages 3 and 6. This leaves mature age 2 males largely unaffected by harvest mortality which results in a small increase in the proportion of age 2 males in the adult population. It has only a small effect on the overall age proportions.

A particularly valuable comparison is between the primary mixed maturation troll harvest scenario (H3) and the terminal fishery counterpart (H5) that adjusts the global harvest rate (hg) downward from the un-restricted MSY rate to the MM Catch Weight Equivalent rate. This achieves approximately the same total catch weight as the mixed maturation troll fishery. This total catch weight (∼8700) is more than 3500 lb smaller than the terminal MSY harvest of 12295 lb (Fig 8). This (H5) terminal harvest scenario catches fewer individuals than the MM fishery (467 versus 728, but since all individuals harvested are mature, the average weight of the fish is significantly larger than the mixed maturation catch (18.5 versus 12.0 lb; Fig 3; Supplementary File S4, Table A). The larger weight of individuals caught in the terminal scenario likely increases their economic value above that of the mixed-stock scenario because larger Chinook may realize a higher price per pound (cf. [37]).

### Differences in total egg deposition between harvest scenarios

For the troll scenarios, the lowest average fecundities occur in the mixed maturation scenario (H3) where the average is 200 eggs per female lower than for either the pre-fishery or the post-fishery averages. For the gillnet scenarios, both the terminal MSY and the mixed maturation MSY scenarios (H2, H4) are more than 350 to 500 eggs per female lower than the pre-fishery averages. Fig 7 shows a related change in the proportion of the average total egg deposition contributed by the two youngest female ages (3 and 4), which increases from an average of 40% in the pre-harvest period to 50% and 52%, respectively in scenarios H3 and H4. This illustrates that the MM harvest scenarios shift the spawner age structure toward younger average ages and renders the population more dependent on egg deposition from the younger, less fecund spawners. In addition, the terminal MSY gillnet scenario (H2) increases the proportion of total egg deposition by age 3 and 4 spawners to 54% due to the strong size selectivity of the gear (Supplement File S4, Table C).

### Model assumptions of note

#### Spawner age and egg size

For both modeling convenience and lack of available data, we assumed that all eggs were the same size and had the same probability of surviving to emergence, and that all emergent parr were of the same size regardless of female age or size, and hence were equal in overall fitness with respect to density dependent survival to the smolt stage. In reality, larger female Chinook deposit larger, better provisioned, eggs that likely possess greater fitness than smaller eggs. As a consequence, the comparisons between the MM fishery scenarios (H3 and H4) and the 4 terminal fishery scenarios may slightly exaggerate the impact of mixed maturation fisheries on population productivity and resilience to the extent that older age spawners are deprived the marginal fitness advantage that may accrue to larger, better provisioned eggs than smaller, younger age females. This is a feature that can be remedied in case-specific applications where population-specific egg-size and/or energy content data are available. However, we believe this effect to be relatively minor compared to the overall change in population structure effected by MM fisheries and thus does not distort the generality of our results.

#### Parameterization of daily growth and density-independent survival rates

Although we lack good data on the actual growth rates associated with maturation at each age, as parameterized in the McGurk growth-mortality equations (Supplementary File S1, A5, equations 5.1 and 5.2), the parameterizations that we chose (Supplementary File S2, Table L) produce reasonable and realistic mean lengths for each of the maturation ages (Fig 2) and realistic age-specific density-independent marine survival rates (Supplementary File S2, Table M). Further, by modeling harvest as length-based, our assumption that older maturing individuals grow more slowly than individuals maturing at younger ages, is more realistic than models that assume that all individuals of a given age in mixed-stock fishery have the same lengths.

#### Fisheries related incidental mortality

An important lacuna in our harvest modeling was ignoring fisheries related incidental mortality (FRIM). FRIM includes retained landings of sub-legal-size fish, post-release mortality of released landed sub-legal, and drop-off mortality of legal and sublegal fish that encounter the gear and die through predation or injury as a consequence. FRIM can be (and normally is) a significant source of fishing mortality in mixed maturation fisheries. We intentionally chose to ignore FRIM in our simulations in order to focus clearly on the impacts to legal size fish. Thus, our results for the 2 MM fishery scenarios are likely an under-estimate of the impact of these fisheries on immature fish. This further highlights the conservative nature of our harvest analyses.

#### Vulnerability of immature Chinook to harvest gear

We assumed that all immature Chinook were equally vulnerable to fishing gear as mature individuals of the same size (fork length). This would require all such immatures to rear in areas where matures are located and where fishing occurs. While likely true for many Pacific Salmon Treaty marine Chinook fisheries in our area of interest, there may be times and areas where this isn’t the case. To the extent to which this is not the case the impact of indirect age-overfishing will be smaller than shown in our model results. Our model, therefore, highlights the most extreme case. To the extent this departs from a specific real-world situation, our results will overestimate the effect of indirect age overfishing. Even in such a case, the harvest of immatures in harvest areas with mature fish will have the kind of detrimental impact on total catch weight, total spawner weight and egg deposition, and age-specific proportions of the catch and spawning population that we show, though the magnitude will be reduced to some degree.

## Conclusion

Our individual-based model of ocean-type Chinook salmon provides for robust evaluations of the likely impacts of different harvest regimes and gear types on the structure and productivity of Pacific coast Chinook populations. It also illuminates the strengths and weaknesses of different harvest regimes for returning benefits to fishers. The results of our modeling exercise provide a succinct analysis of the primary differences between terminal and marine MM fishery regimes on Chinook life-history, productivity, and return to harvesters (measured by catch number and catch weight).

We find that, on balance, terminal fisheries provide greater harvest benefits to fishers than MM fisheries. Additionally, they provide greater protection for the productivity and resilience of the harvested populations. Although we assumed a common egg size independent of female age and size, our results highlight the importance of the greater fecundity of older, larger females to the productivity of Chinook salmon populations, and provide evidence of the importance of monitoring Chinook populations for total egg deposition and the age composition of spawners, rather than (or in addition to) total number of spawners. These results should be of value to scientists and managers concerned with the long-term sustainability of harvestable wild Chinook populations and with the rebuilding of depleted and at-risk populations.

Future planned work examining existing Chinook populations and development of a model applicable to stream-type Chinook, will further our understanding of how recent and past harvest practices have affected wild Chinook populations along the coast of the eastern North Pacific.

The development of appropriately targeted monitoring and assessment projects are also needed to improve understanding of the status of wild Chinook populations at regional and population-specific scales.

## Supporting Information

**S1 File. Chinook survivability parameters and equations. (DOCX)**

**S2 File. Chinook growth parameters. (DOCX)**

**S3 File. Characteristics of the equilibrium spawner abundance of the IBDEM Chinook model. (DOCX)**

**S4 File. Tables of harvest simulation results. (DOCX)**

**S5 Table. Summary of a representative 1000 year simulation. (XLSX)**

## Acknowledgments

We thank Marco Castellani for generously sharing all of the code for the IBSEM model [38] and for answering questions regarding some aspects of the coding of the model. We thank Jeffrey Hard (Conservation Biology Division, Northwest Fisheries Science Center, National Marine Fisheries Service, retired) for reviewing the coding of the maturation genetics in an earlier version of the model and for providing advice and answering questions regarding the expected magnitude of the heritability of maturation. We are grateful to Marty Kardos of the Conservation Biology Division, Northwest Fisheries Science Center and Robin Waples of the Northwest Fisheries Science Center (retired) for providing helpful comments on a draft of the current paper, particularly our approach to coding the quantitative genetics. These all greatly improved the organization of the paper. This does not, however, imply any endorsement of our approach and results by these reviewers or NMFS.

